# SARS-CoV-2 Spike H655Y drives protease preference but does not dictate cellular tropism

**DOI:** 10.64898/2026.07.05.736580

**Authors:** Diego Cantoni, Wilhelm Furnon, Sarah Little, Vanessa Cowton, Bridget Larman, Reehana Damasceno, Brian Willet, Arvind H. Patel, Massimo Palmarini, Joe Grove

## Abstract

Throughout the COVID-19 pandemic, SARS-CoV-2 has undergone rapid adaptation, with the Omicron variant exhibiting an unexpected shift towards upper airway infection and preference for membrane fusion activated by endosomal cathepsins, a reversion to ancestral sarbecovirus entry mechanisms. This phenotype coincides with convergent acquisition of the spike mutation H655Y, which reduces TMPRSS2-mediated activation. Here, using a comprehensive panel of 13 spikes of SARS-CoV-2 variants, we interrogated whether H655Y-mediated protease switching explains Omicron’s upper airway phenotype, challenging several existing hypotheses, from spike stability, shedding and acquired intra-molecular interactions by H655Y. Our findings reveal that protease preference and tissue tropism are mechanistically uncoupled. While spike pre-processing by furin determines protease preference in pre-Omicron variants, this relationship breaks down in H655Y-bearing viruses. Notably, mutations in the NTD and RBD of BA.2.86 can override the H655Y phenotype entirely, indicating that RBD-mediated interactions, rather than protease usage, represent the critical determinants of Omicron’s upper airway adaptation. This work reframes our understanding of coronavirus spike evolution, revealing that tissue tropism operates through mechanisms fundamentally distinct from those dictating protease preference.

## Introduction

Throughout the COVID-19 pandemic, severe acute respiratory syndrome coronavirus 2 (SARS-CoV-2) faced complex selective pressures including, initially, adaptation to a novel human host and with time evolving population immunity, driven by natural infection and vaccination. As a consequence, the spike protein, amongst other viral proteins, has undergone continuous evolution, leading to the emergence of variants defined by constellations of mutations which work cooperatively to modulate spike stability, receptor affinity and tissue tropism, and to mediate immune escape [1–4].

The trimeric spike protein complex is the key mediator for virus attachment and entry [5,6]. Each monomer of the spike is composed of two subunits, S1 and S2. The S1 subunit contains the N-terminal domain (NTD) and the receptor binding domain (RBD), which binds to ACE-2 receptor. The S2 subunit contains the class-I fusion machinery required for membrane fusion. The S1/S2 boundary of SARS-CoV-2 spike is unique across all known sarbecoviruses in that it contains a multibasic cleavage site [7]. Upon spike biogenesis, this is proteolytically cleaved by the trans-Golgi network-resident convertase furin; hence this site is commonly referred to as the furin cleavage site (FCS). S1/S2 cleavage destabilises spike and is reported to increase entry efficiency as more RBDs on the trimer are in the conformationally exposed ‘up’ position, priming it for attachment [8,9]. It also promotes the rapid shedding of S1 after further proteolytic cleavage by TMPRSS2 at the S2’ site, which is located downstream of the FCS and is necessary to liberate the fusion peptide for insertion into target cellular membranes. Subsequent conformational change within S2 pulls the viral and cellular membrane together, driving their fusion, and completing the entry phase of infection [10]. By comparison, SARS-CoV-1 spike protein does not possess a FCS at the S1/S2 junction, and whilst it can be processed by TMPRSS2 at the cell surface [11], the spike cleavage is delayed and often occurs after endocytosis, following processing by endosomal cathepsins [12,13].

In the early stages of the pandemic, the preferred tropism of SARS-CoV-2 was the lower airway [14,15], exploiting efficient TMPRSS2-mediated proteolysis, to achieve fusion from the plasma membrane. This phenotype was largely attributed to the FCS, as the spike is already ‘primed’ for fusion as it arrives at the cell surface [16]. However, the Omicron BA.1 and BA.2 variants, which possess heavily mutated spike proteins, not only exhibited extensive immune evasion but also altered tissue tropism, with a preference for replication in the upper-airway [17,18]. Following their emergence, BA.1 and BA.2 underwent explosive transmission, associated with vaccine escape but relatively mild illness. This represents a defining moment in the pandemic with all subsequent major SARS-CoV-2 lineages stemming from the BA.2 Omicron event.

These changes in tissue tropism, pathogenesis and immune escape were accompanied by phenotypic changes in spike, as measured in various *in vitro* assays. In particular, in cells lines, Omicron lineage viruses exhibited a switch from TMPRSS2-mediated fusion at the cell surface to cathepsin-dependent fusion from the endosome; akin to the delayed fusion process of SARS-CoV, despite the preservation of the FCS within the Omicron spike [17,18]. Studies have reported mutations within the FCS, proximal to the FCS and in the RBD of spike as potential molecular determinants of this phenotype [19–22]. Whilst studies in lung-derived human airway organoids suggest that Omicron retains TMPRSS2 entry in lower respiratory tissue [23], it has been suggested that a preference for endosomal fusion may determine enhanced replication in the upper-airway [24–28]. Although others ascribe this to acquired sensitivity towards cell surface metalloproteases [29,30], and cellular sialoglycans [31], or to differential usage of subpopulations of ACE2 receptor [32].

We aimed to delineate the molecular determinants of protease preference and, critically, how this contributes to tissue tropism. To achieve this, we performed a systematic comparison of spike protein phenotypes from throughout the pandemic, finding that the common attribute of cathepsin-using spikes was the H655Y mutation. We demonstrate that H655Y has opposite effects on cathepsin and TMPRSS2-dependent entry, rescuing one and limiting the other. Testing various mechanistic hypotheses, including spike stability and newly acquired intra-molecular interactions, revealed them insufficient to explain the phenotype. Ultimately, using primary human airway cells, we uncovered that protease preference and tissue tropism are mechanistically uncoupled, implicating RBD-mediated interactions as critical drivers of Omicron’s upper airway adaptation.

## Materials and Methods

### Cell Culture

HEK293T cells, HEK293T CD81-KO cells [33] and A549-ACE2-TMPRSS2 (AAT) [34] were cultured using Dulbecco’s Modified Eagle Medium (DMEM) (Thermofisher Scientific) supplemented with 10% foetal bovine serum (FBS) (Thermo fisher Scientific), 1% penicillin/streptomycin (Thermofisher Scientific) and 1% non-essential amino acids (Thermofisher Scientific). Calu3 cells were maintained in DMEM culture medium supplemented with 20% FBS and 1% penicillin/streptomycin. Cells were routinely passaged to avoid overconfluence and maintained at 37 °C and 5% CO_2_ in a humidified chamber. AAT cells were used for a maximum of 20 passages. All cells were verified to be free of mycoplasma.

Primary human reconstituted nasal epithelial cells (hNEC) were purchased from Epithelix and maintained in MucilAir complete culture medium at an air-liquid interface.

### Plasmids, Spike Mutagenesis and Domain Swap Constructs

Plasmids containing codon-optimised SARS-CoV-2 spike genes, A.30, Gamma (hereafter referred to as P.1), BA.1, BA.2 and BA.2.86 were designed by GeneArt, Thermofisher, and inserted into a pcDNA3.1+ vector. The following variants were gifted by the G2P-UK consortium: Wu-Hu-1, D614G, Alpha, Beta, Delta, BA.4/5, and XBB. All SARS-CoV-2 spikes contain an 18-amino acid C-terminus deletion. Pangolin spike (isolate cDNA18-S) was inserted into a pD603 expression plasmid.

To generate domain swapped spike plasmids, we took advantage of restriction digest sites situated between the NTD, RBD, FCS and S2 domains. For mutagenesis, fragments containing the desired mutations were ordered via Twist Biosciences, restriction digested and ligated using Quick-ligase (New England Biolabs). All sequences were verified by either Sanger sequencing (SourceBiosciences) or by whole plasmid sequencing (Plasmidsaurus).

For N-terminus HA-tagging of spikes, fragments containing HA-tags after the signal peptides of the spike were ordered via Twist Biosciences, and inserted using restriction digestion and ligation. All sequences were verified by Sanger sequencing.

### Pseudotype Virus Production

HEK293T CD81-KO cells were used to produce high titres of SARS-CoV-2 PVs, as previously described by Kalemera et al. [33]. Briefly, Cells were seeded in 6-well plates at a density of 1×10^6^ cells per well and transfected the following day with 1.3μg p8.91 lentiviral packaging plasmid, 1.3μg pCSFLW luciferase reporter and 200ng of spike expression plasmid, using Fugene 6 (Promega) at a 3:1 ratio according to the manufacturer’s protocol. Transfection complexes were incubated for 15 minutes before adding transfection complexes onto the cells.

Supernatants were harvested at 48 hours and 72 hours post transfection and passed through at 0.45μm cellulose acetate filter prior to use or storage at −20 °C.

To inhibit furin pre-processing during PV production, HEK293T CD81-KO cells culture medium was removed and replenished 4 hours post transfection with fresh media containing 60μM of furin inhibitor Decanoyl-RVKR-CMK (CMK) (Sigma-Aldrich). An equal volume of DMSO was used as a negative control. Supernatants were harvested at 48 and 72 hours post-transfection and passed through at 0.45μm cellulose acetate filter prior to use or storage at −20 °C.

### Pseudotype Virus Infection Assays

To evaluate PV entry, HEK-293T, AAT or Calu3 cells were seeded in white 96-well F-bottom plates at a density of 30,000 cells per well, followed by addition of equal volumes of undiluted PV. Infections were performed with a minimum of 4 technical replicates and 3 biological replicates.

For viral entry drug inhibitor assays, AAT cells were treated with 10μM of either camostat mesylate (Sigma-Aldrich) or E64d (Sigma-Aldrich) for 2 hours prior to infection. An equal volume of DMSO was used as a negative control.

For the ACE2 and TMPRSS2 usage infection assay, HEK-293T cells in 6-well plates were transfected with either 600ng of ACE-2 plasmid, 50ng of TMPRSS2 plasmid, both plasmids or an empty vector plasmid as a control transfection. The following day, the transfected cells were trypsinised and seeded as described above prior to infection. For this assay, cells were infected with ten-fold serial dilutions of PV in DMEM, from 1:2 to 1:20,000.

All infections were incubated for 48 hours prior to lysis using 25μL neat Bright-Glo luciferase reagent (Promega). Luciferase reporter activity was measured using a Glo-Max luminometer (Promega).

### Generation of recombinant viruses using reverse genetics

Recombinant viruses were generated using transformation-associated recombination (TAR) in yeast (*Saccharomyces cerevisiae*). Briefly, bacterial artificial chromosome vectors (BAC) containing the full-length genome of SARS-Cov-2 were cut by restriction enzymes and combined with overlapping fragments containing the desired single-point mutation in spike (H665Y/Y655H). The fragments were assembled in yeast using TAR and DNA was isolated from single-yeast clones. The yeast DNA was transformed into *E.coli* and again DNA was purified from single colonies to yield BAC vectors carrying the H655Y/Y655H mutation swap. Recombinant virus was rescued following direct transfection of BAC vector DNA into IGROV-1 cells.

### Virus propagation and titration

Working virus stocks were generated and titrated as previously described. In brief, 50–100 μl of P0 virus stock was inoculated onto Vero E6 cells (5 × 10^6 cells seeded in a T-75 flask). Following a 1 h incubation period, the inoculum was removed and replaced with DMEM supplemented with 2% FBS and 1% penicillin–streptomycin. Culture supernatants were harvested upon the appearance of cytopathic effect (CPE), typically observed between 4 and 6 days post-infection, and clarified by centrifugation at 500 × g before storage at −80 °C. Virus stock titres were determined by reverse transcription quantitative PCR (RT–qPCR) and expressed as Orf1a gene copy number-equivalent genomes per millilitre of supernatant.

Two clones of each rescued virus mutant were passaged in Vero E6 cells to produce P1 working stocks for use in replication assays. Viral RNA was extracted using a QIAamp viral RNA mini kit (Qiagen, 52906) and viral genome sequenced using Oxford Nanopore as previously described [35].

### Virus replication assays

Calu-3 cells were seeded in 24-well plates at a density of 1.8 × 10^5 cells per well and infected 5 days later with 10^4 Orf1a genome copy equivalents per well in serum-free RPMI-1640 medium. Following a 1 h incubation at 37 °C, the inoculum was removed, the cells were washed, and fresh RPMI-1640 medium supplemented with 20% FBS was added. Culture supernatants were collected at the indicated timepoints, and viral RNA was extracted and quantified as described above.

For primary hNECs, cultures were washed once with serum-free DMEM before apical infection with 10^5 Orf1a genome copy equivalents per well in serum-free medium. Cells were incubated at 37 °C in a humidified atmosphere containing 5% CO₂ for 2 h, after which the inoculum was removed and the cultures were washed once with serum-free DMEM. The first sample was collected immediately following this wash. At the indicated timepoints, virus released from the apical surface was harvested by adding 100 µl of pre-warmed serum-free DMEM to the apical compartment and collecting the supernatant after a 15 min incubation at 37 °C. Samples were subsequently mixed with LBF lysis buffer (Beckman Coulter, A35604) and heated at 56 °C for 20 min for complete inactivation.

### RNA extraction and RT-qPCR

Viral RNA was extracted from culture supernatants using the RNAdvance Blood Kit (Beckman Coulter, A35604) according to the manufacturer’s instructions. Extracted RNA was subsequently used as a template for the detection and quantification of viral genomes by reverse transcription quantitative PCR (RT–qPCR) using the Luna Universal Probe One-Step RT–qPCR Kit (New England Biolabs, E3006E). SARS-CoV-2 RNA was specifically detected by targeting the ORF1a gene with the following primers and probe: SARS-CoV-2_Orf1a_Forward, 5′-GACATAGAAGTTACTGGCGATAG-3′; SARS-CoV-2_Orf1a_Reverse, 5′-TTAATATGACGCGCACTACAG-3′; and SARS-CoV-2_Orf1a_Probe, ACCCCGTGACCTTGGTGCTTGT, containing HEX/ZEN/3IABkFQ modifications. A SARS-CoV-2 RNA standard curve was used in each run, enabling quantification of viral genomes, which were expressed as the number of Orf1a RNA molecules per millilitre of supernatant. All RT–qPCR assays were performed on an ABI7500 Fast instrument, and data were analysed using 7500 Software v.2.3 (Applied Biosystems, Life Technologies).

### Western Blotting

Transfected cells were washed with ice-cold PBS and lysed with lysis buffer containing 50 mM Tris/HCl pH 7.5, 1 mM EGTA, 1 mM EDTA, 1% (v/v) Triton X-100, 1 mM sodium ortho-vanadate, 50 mM sodium fluoride, 5 mM sodium pyrophosphate, 0.27 M sucrose, 10 mM sodium 2-glycerophosphate, 1 mM phenylmethylsulphonyl fluoride, 1 mM benzamidine and 1X NuPAGE LDS sample buffer (Thermo Fisher Scientific) [36]. PVs were purified from supernatant using Dynabeads Intact Virus Enrichment (Thermo Fisher Scientific) kit according to the manufacturer’s protocol until the last step where 60μL lysis buffer was directly added to the beads containing enriched PVs. Samples underwent SDS-PAGE using pre-cast 4-20% Mini-PROTEAN TGX precast gels (Bio-Rad), followed by transfer to a PVDF membrane using a Trans-Blot Turbo transfer system (Bio-Rad). Membranes were blocked with 5% milk in PBS + 0.1% Tween-20 (PBS-T) and then probed with primary antibodies overnight at 4 °C: mouse anti-spike S2 (Genetex GTX632604), mouse anti-spike S1 (Cell Signalling E7M5X), rabbit anti-p55+p24+p19 (Abcam 63917), rabbit anti-HA-tag (Cell Signalling C29F4) diluted in blocking buffer overnight at 4 °C. Membranes were washed 4 times with PBS-T and incubated with secondary antibodies in blocking buffer for 1 hour at room temperature: anti-rabbit IgG (H + L) DyLight 800 conjugate (Cell Signalling 5151S), anti-mouse IgG (H + L) DyLight 680 conjugate (Cell Signalling 5470S), goat anti-mouse IgG (H + L) cross absorbed HRP conjugate (Invitrogen G-21040), β-actin-HRP conjugate (Thermo MA5-15739-HRP). Membranes were washed 4 times prior to visualisation with Odyssey CLx imager (Li-Cor) or by enhanced chemiluminescence (ECL) (Pierce) with Chemidoc MP (Bio-Rad). Image analysis was performed using Image Studio Lite Software (Li-Cor).

### Phylogenetic Analysis

The SARS-CoV-2 whole genome phylogenetic tree containing 7,725,737 sequences from International Nucleotide Sequence Database Collaboration (INSDC databases) was downloaded from Taxonium.org on the 1^st^ June, 2026, with position 655 annoted on the web server [37,38]. For sarbecovirus spike tree, we downloaded 526 sarbecovirus spike amino acid sequences from NCBI, and aligned using MAFF-T L-INS-i settings. Sequences that showed poor alignment or large gaps were manually deleted. The phylogenetic tree was constructed using IQ-TREE2 [39] on the trimmed and aligned dataset. Branch support was assessed using 1,000 ultrafast bootstrap replicates and the best model was selected using ModelFinder integrated in IQ-TREE2.

### Statistical Analysis

All statistical analyses were carried out using GraphPad Prism v.10.2.2. Unless stated otherwise, the results are expressed as mean ± s.e.m. Data distribution was assessed by Kolmogorov-Smirnov test to determine normality and inform on subsequent statistical tests (data not shown).

## Results

### Convergent acquisition of spike:H655Y caused switches in SARS-CoV-2 protease usage on multiple occasions throughout the pandemic

To capture the evolutionary trajectory of SARS-CoV-2 we examined the main Variants of Concern (VOCs), designated by the World Health Organisation (WHO), as well as variant under monitoring, A.30, and two ancestral relatives, SARS-CoV and Pangolin-CoV. We performed pseudotype virus (PV) entry assays in A549-ACE2-TMPRSS2 (AAT) cells which permit both surface entry and endosomal entry. This allowed us to modulate entry pathway availability using two pharmacological inhibitors: camostat (blocks TMPRSS2/cell surface fusion) and E64d (blocks cathepsin/endosomal fusion). We found that ancestral relatives SARS-CoV-1 and Pangolin-CoV utilise the endosomal route most efficiently, consistent with previous studies [40,41] (Figure 1A). We also confirmed that most VOCs prior to BA.1 preferentially utilise TMPRSS2 to achieve cell surface entry. However, the P.1 VOC and A.30 variant appeared to be sensitive to both TMPRSS2 and cathepsin. This suggests that these spikes do not have a strict dependence towards either protease. For BA.1 through to XBB, we observed a switch towards cathepsin dependency, a hallmark of the Omicron lineage. However, the later BA.2.86 variant appears to revert towards TMPRSS2 usage which is consistent with our recent finding usings authentic virus [42].

**Figure 1:**
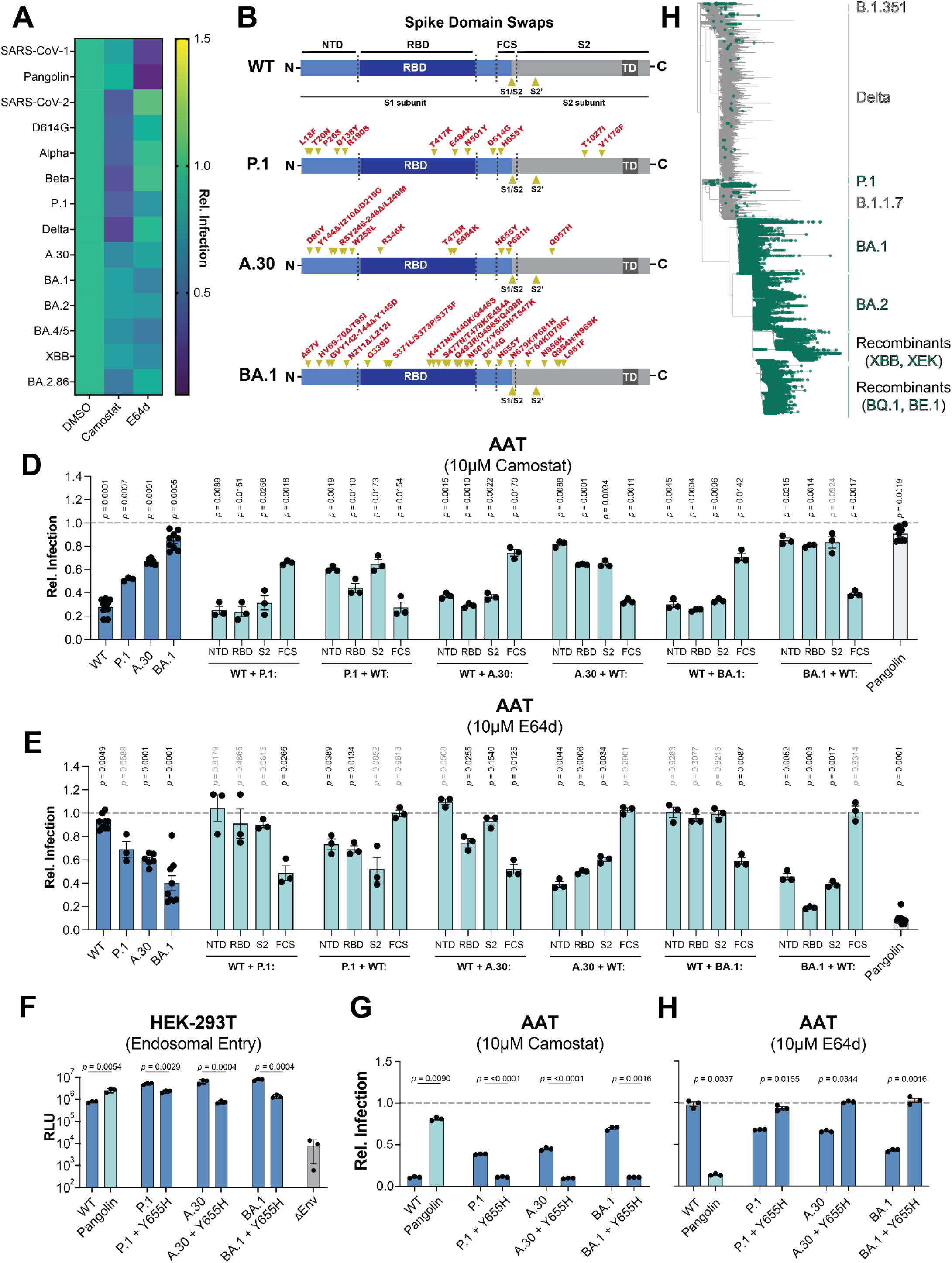
Spike determinants for enzyme usage. **A.** Heat map of AAT cell entry assay depicting relative PV infections of SARS-CoV-2 variants in the presence of 10μM camostat or E64d, with DMSO as a control. **B.** Ribbon diagrams depicting the domain swapped spike designs and mutational profiles of P.1, A.30 and BA.1 spikes. **C, D**. Sensitivity panel of domain swapped spike PVs in AAT cells in the presence of 10μM camostat or 10μM E64d. The dotted line represents relative viral infection with DMSO control. **E.** Infectivity of variant and reverted Y655H mutant PVs in TMPRSS2-deficient HEK293T cells. dEnv is bald PV control. **F, G.** Sensitivity panel of Y655H reverted spike PVs in AATs in the presence of 10μM camostat or 10μM E64d. The dotted line represents relative viral infection with DMSO control. **H.** Phylogenetic tree of SARS-CoV-2 taken from Taxonium.org on 01/06/2026. Green dots represent sequences that contain H655Y. For **C, D,** and **E**, statistical significance was determined by one-sample *t*-test. For **F,** statistical significance was determined by *Welch’s t*-test. *P* values ≤ 0.05 are considered significant. A minimum of n=3 was used for each comparison.

Given the evidence for changes in protease preference in both pre- (P.1, A.30) and post-Omicron (e.g. BA.1) viruses, we next sought to determine which regions of the spike govern this preference. We performed reciprocal domain swaps of NTDs, RBDs, S2, and a region spanning subdomain 2 (SD2, which is proximal to the FCS) and FCS from variants P.1, A.30 and BA.1 against WT (Wu-Hu-1) spike (Figure 1B). Using the same inhibitor assay in AAT cells, we observed that in all three backgrounds, switches in protease preference correlate with swapping the SD2/FCS region (Figure 1C & D). In swaps where WT virus received a SD2-FCS from P.1, A.30 or BA.1, we observed increased sensitivity to E64d and reduced inhibition by camostat, indicating a preference for cathepsin usage. Conversely, in reciprocal mutations where P.1, A.30 and BA.1 received a WT SD2-FCS we observed an increased sensitivity to camostat and a complete loss of inhibition by E64d, indicating a preference for TMPRSS2 usage (Figure 1C & D). Each of these spike variants have different mutational profiles and the only mutation shared by all three is H655Y, which sits in the SD2/FCS region. We, therefore, hypothesised that this mutation is a master determinant of protease preference.

To confirm the importance of H655Y for capthepsin-dependent endosomal entry we infected HEK293T cells, which do not express TMPRSS2 and therefore only support cathepsin-mediated endosomal entry. For each endosomal preferring spike (P.1, A.30, and BA.1), the singular amino acid reversion (Y655H) was sufficient to decrease HEK293T infectivity to levels comparable to that of WT spike (Figure 1E). Moreover, protease switching of single-residue revertants was also confirmed via infection of AAT cells infections in the presence of camostat (Figure 1F) and E64d (Figure 1G). These data show H655Y alone is sufficient to explain preference for activating protease and subcellular site of membrane fusion: cell surface or endosome. This mutation has been acquired on multiple independent lineages throughout the SARS-CoV-2 phylogeny (Figure 1H), including two branches leading to VOCs (P.1 and Omicron). Therefore, our observations suggest that SARS-CoV-2 explored changes in protease preference throughout the pandemic.

### Spike pre-processing by furin determines protease preference in pre-Omicron variants, but not in Omicron and H655Y bearing viruses

It has been hypothesised that H655Y is a stabilising mutation that promotes integrity of the spike S1:S2 complex following furin cleavage [43]. By extension, H655Y may represent a specific adaptation to the FCS, which remains a unique feature of SARS-CoV-2 compared to other sarbecoviruses. To delineate the intrinsic effects of possessing a multibasic cleavage site at the S1/S2 boundary, from the inherent destabilising effects of furin processing during spike biogenesis, we used the furin inhibitor CMK.

Infection of HEK293T cells (which only support cathepsin-dependent endosomal entry), Calu3 cells (which only support TMPRSS2-dependent cell surface entry) and AAT cells (which support both modes) with WT virus produced under increasing concentrations of CMK revealed distinct effects (Figure 2A). CMK treatment uniformly enhanced entry into HEK293T cells by ∼10-fold. This suggests that a majority of potential endosomal entry events fail under normal conditions, but can be “rescued” by preventing pre-processing of spike; this is consistent with stability being important for this entry route.

**Figure 2:**
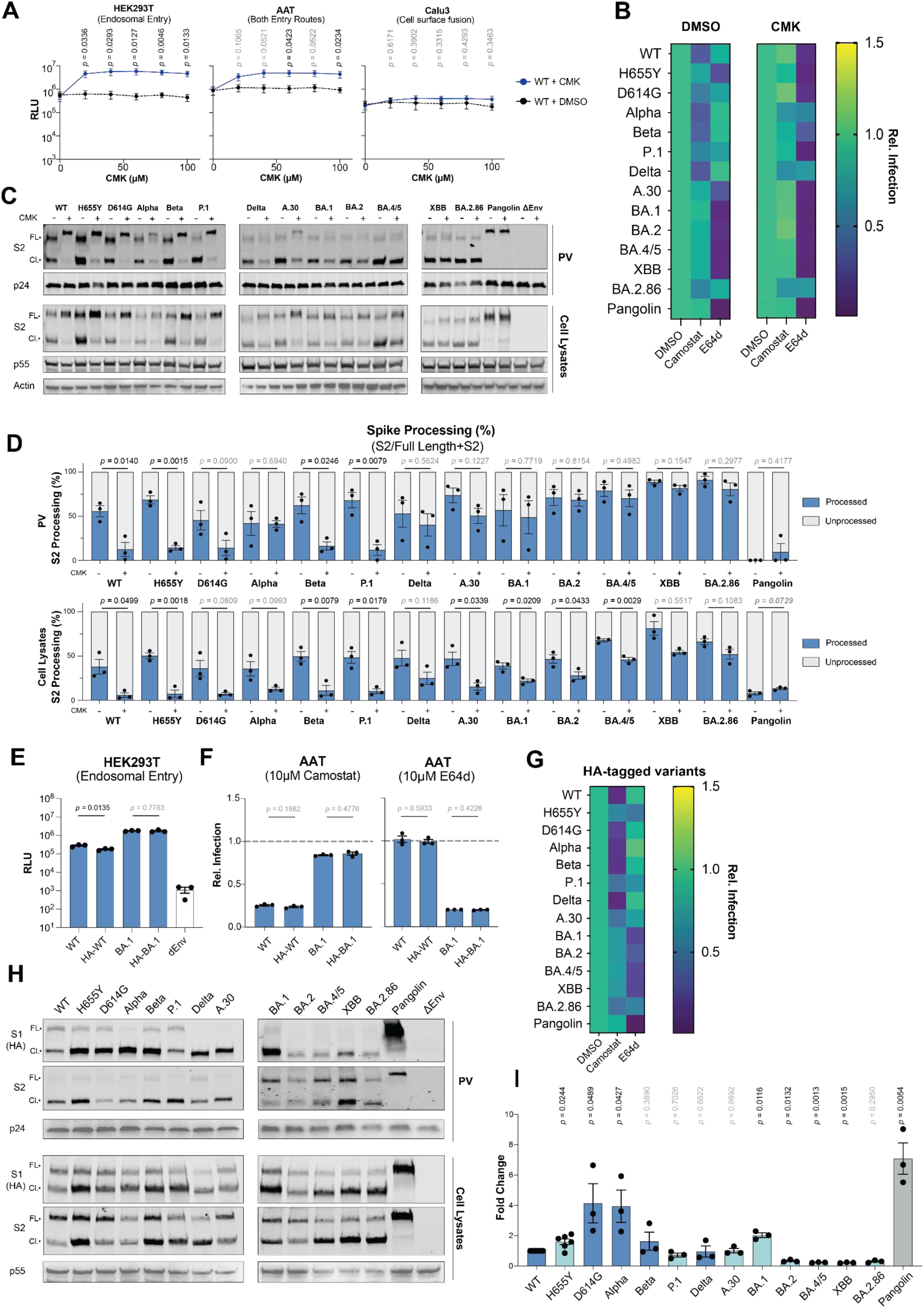
H655Y characteristics on spike cleavage and stability. **A.** Infection of HEK293T, ATT or Calu3 cells with WT PVs produced in the presence of increasing concentrations of CMK. **B.** Heat map of AAT cell entry assay depicting relative PV infections of SARS-CoV-2 variants produced in the presence of either DMSO or 60μM CMK in AAT cells treated with 10μM camostat, E64d or DMSO as a control **C** Western blot panels of SARS-CoV-2 spike variants produced in presence of either DMSO or 60μM CMK. Lysates and PVs were immunoblotted against S2 (spike, FL: full length, Cl: cleaved/processed), p55/p24 (gag-pol control) and actin. **D**. Quantification of spike processing from western blot images. **E.** PV infections of HA-tagged and non-tagged spikes in HEK293T cells. **F.** Sensitivity panel of HA-tagged and non-tagged PVs in AATs in the presence of 10μM camostat or 10μM E64d. The dotted line represents relative viral infection with DMSO control. **G.** Heat map of AAT cell entry assay depicting relative PV infections of HA-tagged SARS-CoV-2 variants in the presence of 10μM camostat or E64d, with DMSO as a control. **H.** Western blot panels of HA-tagged spikes. Lysates and PVs were immunoblotted against HA-tag (spike S1), S2 (spike S2), p55/p24 (gag-pol control) and actin. **I.** Bar chart quantifications of total S1 relative to total S2, calculated from three independent western blots of purified PV. For **A, D, E** and **F,** statistical significance was determined by *Welch’s t*-test. For **I,** statistical significance was determined by one-sample *t*-test. *P* values ≤ 0.05 are considered significant. A minimum of n=3 was used for each comparison.

Previous studies have demonstrated reduced infection of Calu3 cells by FCS-deleted viruses [16,44]. However, in our experiments CMK had no effect on WT spike-mediated entry into this cell type. This indicates that whilst the multibasic FCS may be necessary for efficient TMPRSS2-dependent entry, furin cleavage *per se* is not required. This agrees with other reports [45], and suggests that in the absence of pre-processing, cleavage of the multibasic S1/S2 boundary can occur efficiently during entry into Calu3 cells. Entry into AAT cells exhibited an intermediate profile with moderate enhancement of entry, consistent with their supporting both pathways. In summary, furin pre-processing is detrimental to endosomal entry and this may have provided a selection pressure for adaptation.

Next, we explored the extent of furin pre-processing in our panel of variants, plus the H655Y mutant, and investigated how this impacts protease preference (Figure 2B, C & D). To confirm that CMK treatment inhibited furin pre-processing during PV biogenesis, we analysed both purified PVs and cell lysates by western blot (Figure 2C). The majority of pre-Omicron spikes exhibited reduced cleavage upon CMK treatment, though Alpha and Delta variants were notable exceptions. In contrast, none of the post-Omicron spikes and the phenotypically related A.30 spike displayed defects in spike cleavage. Furthermore, the spikes from BA.1 onwards showed the highest pre-processing efficiencies across all variants in our panel, both with and without CMK treatment (Figure 2D). This suggests that H655Y, present in these spikes, does not negatively modulate furin cleavage of spike. An implication of this result is that Omicron lineage pre-processing may occur in a furin-independent manner.

To examine whether pre-processing defects alters activating protease preference, we infected AAT cells with CMK +/− treated PVs, and using camostat and E64d to differentiate entry pathway (Figure 2B). Those pre-Omicron spikes that were sensitive to CMK treatment exhibited a dramatic shift to endosomal entry, as evidenced by potent inhibition by E64d. Alpha and Delta, had a smaller shift in protease usage, concomitant with the absence of pre-processing defects in these spikes (Figure 2C & D). Variant P.1 also had a less pronounced shift, owing to its increased usage of endosomal entry as demonstrated in Figure 1. By contrast, CMK treatment had little to no effect on the protease preference of H655Y, A.30, and the Omicron variants.

In summary, inhibition of furin pre-processing redirects pre-Omicron variants towards endosomal entry and the extent of this switch correlates with the extent of pre-processing inhibition. Spikes possessing H655Y, either alone or in a wider mutational background, facilitate entry via the endosomal route, irrespective of CMK treatment or pre-processing status. The notable exception is BA.2.86, which once again displayed a more pre-Omicron-like phenotype, with and without CMK. These data are consistent with the hypothesis that H655Y is a compensatory adaptation to the FCS, potentially overcoming the deleterious effects of spike pre-processing on endosomal entry.

### S1 shedding does not correlate with residue 655 genotype

Given that the effect of CMK treatment on protease usage, broadly, mirrors that of H655Y, we hypothesised that H655Y may be stabilising the S1:S2 complex following furin pre-processing. This property can be assessed by quantifying the amount of the sheddable S1 subunit remaining associated with virus particles. Several studies have tested levels of S1 using polyclonal antibodies. However, to ensure consistent detection across our broad panel of variants, which contain heavily mutated S1 subunits, we opted to N-terminally tag spike. Control infections confirmed that the N-terminal HA-tag minimally interferes with virus production (Figure 2E), and maintains the entry phenotype observed by the variants throughout the study (Figures 2F, 2G). We quantified S1 and S2 from both purified PVs and cell lysates by western blotting. To account for slight differences in spike incorporation between variants, we calculated S1 levels relative to S2 for each variant (Figure 2H & I). As S2 is anchored to the viral membrane, it serves as a readout of spike incorporation for each PV.

Pangolin-CoV acts as a control as it lacks a multibasic cleavage site and thus S1 remains covalently bound until virus entry. Accordingly, S1 levels for Pangolin-CoV spike were consistently > 5-fold greater than WT SARS-CoV-2. If H655Y promotes S1:S2, stability we would expect greater S1 retention in spikes possessing this mutation, akin to the result with Pangolin-CoV. While we observed a 2-fold increase in S1 for WT+H655Y compared to WT only, this correlation between 655 genotypes and S1 levels was not maintained across the panel of variants. For example, the D614G and Alpha variant spike, which do not possess the H655Y mutation, retained high levels of S1, whereas the latter Omicron variants, which do possess H655Y, appeared to more-readily shed S1. Therefore, the entry phenotype of H655Y bearing viruses cannot be simply ascribed to spike complex integrity.

### H655Y reduces the ability of spike to undergo TMPRSS2-mediated activation

Given that our inhibitor assays with AAT cells reveal a switch in protease preference, we wanted to better understand if this represented a rescue of cathepsin-mediated entry (similar to that seen in CMK treated virus), or whether TMPRSS2 priming of spike was directly impacted. To examine this, we performed PV infections of HEK293T cells exogenously expressing TMPRSS2 alone, ACE2 alone, or both ACE2 and TMPRSS2, thereby recapitulating the cell surface fusion apparatus in cells that otherwise support only cathepsin-mediated endosomal entry.

In initial experiments, we infected cells with serial dilutions of PVs bearing WT, H655Y, BA.1 and BA.1 revertant Y655H spikes (Figure 3A). Considering WT spike alone, we observed two features. First, exogenous TMPRSS2 expression alone did not enhance infection. This suggests that ACE-2 levels are limiting in unmodified HEK293T cells and is consistent with a two-step process where TMPRSS2 processing follows receptor engagement (Figure 3A). Indeed, co-transfection of ACE-2 unlocks the enhancing capacity of TMPRSS2. Second, at low PV dilutions (e.g. 1/2) signal saturates, masking the relative differences between variants. Therefore, we focussed subsequent comparisons on 1/200 diluted PV, which sits within the linear range of the assay.

**Figure 3:**
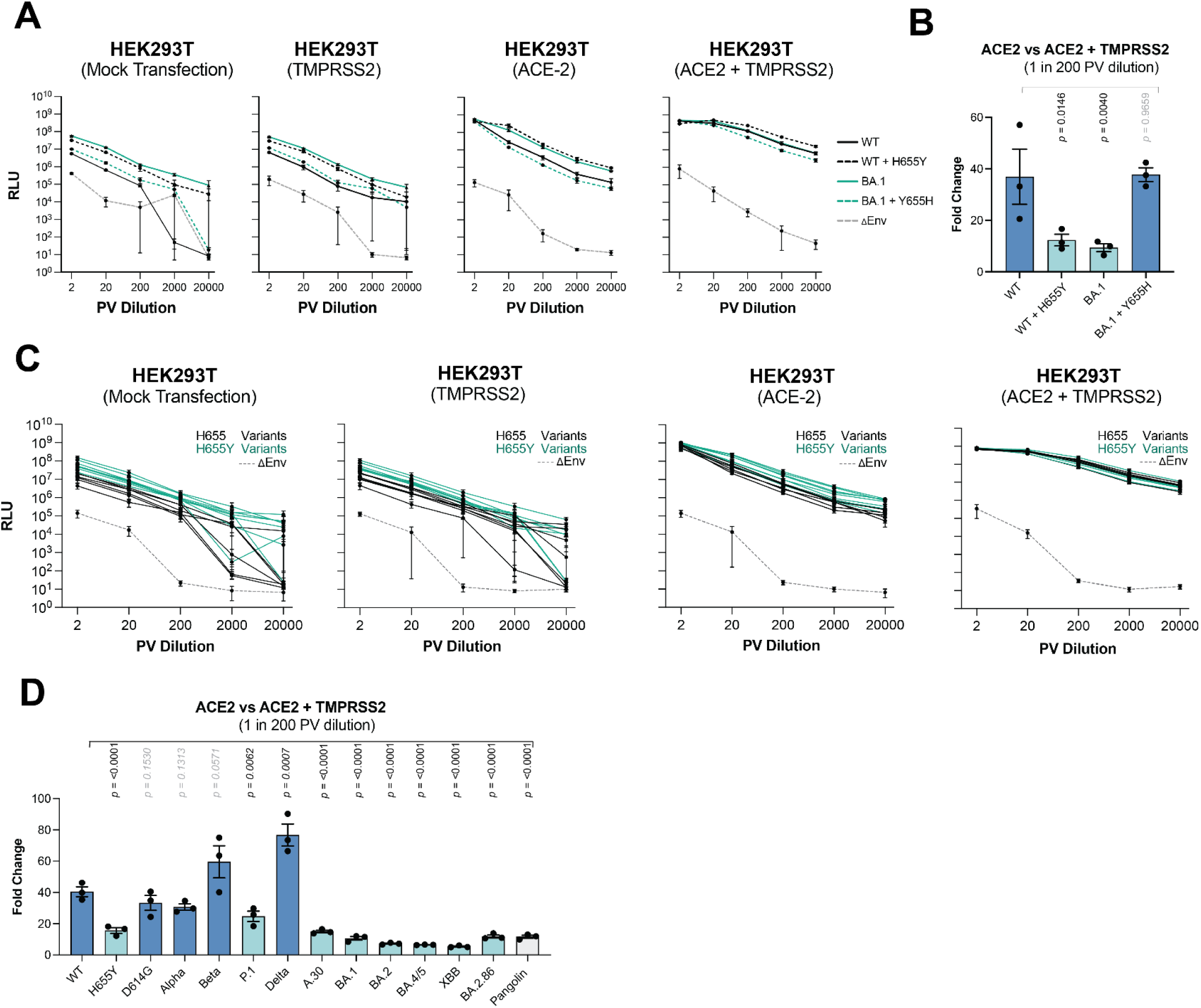
Altered TMPRSS2 usage in H655Y containing spikes. **A.** Serial dilution PV infections in HEK293T cells transfected with either empty vector, TMPRSS2, ACE2 or ACE2+TMPRSS2. **B.** Fold change in PV infectivity at 1:200 serial dilution between ACE2 only and ACE2+TMPRSS2 conditions. Dark blue colour represents spike with H655, teal represents spike with Y655. **C.** Serial dilution PV infections of all variants in HEK293Ts transfected with either empty vector, TMPRSS2, ACE2 or ACE2+TMPRSS2. **D.** Fold change in PV infectivity of all variants at 1:200 serial dilution between ACE2 only and ACE2+TMPRSS2 conditions. Dark blue colour represents spikes with H655, teal represents spikes with Y655. For **B** and **D,** statistical significance was determined by one-way ANOVA test was used against the mean of WT. *P* values ≤ 0.05 are considered significant. A minimum of n=3 was used for each comparison.

By comparing PV infection in HEK 293T cells expressing exogenous ACE-2 to those expressing both ACE-2 and TMPRSS2, we can assess the ability of a given virus to undergo TMPRSS2-mediated activation. Under these conditions, WT virus infection was enhanced by ∼40-fold suggesting efficient use of TMPRSS2. By contrast, H655Y and BA.1 exhibited only a ∼10-fold increase (Figure 3B). Moreover, the BA1 revertant Y655H was restored to a WT-like phenotype, with a ∼40-fold increase. We have already established that H655Y rescues cathepsin-mediated endosomal entry (Figure 2). This experiment goes further by suggesting that this mutation also reduces the ability of SARS-CoV-2 to undergo TMPRSS2-mediated cell surface entry.

To consolidate this finding, we extended the study to all SARS-CoV-2 variants shown in Figure 1. In unmodified HEK293T cells or those singly expressing exogenous ACE2 or TMPRSS2, variants containing H655Y outperformed all H655 variants (Figure 3C). However, when ACE2 and TMPRSS2 are co-expressed we found that H655 variants experience a strong TMPRSS2 boost, such that all viruses exhibited similar infectivity levels. Comparison at 1/200 PV dilution reveals a clear pattern; infectivity of H655 containing variants was strongly enhanced by the presence of TMPRSS2. Most notably, Beta and Delta variants exhibited a 60-80-fold boost in infection. In contrast, H655Y containing variants were only moderately enhanced in the presence of TMPRSS2, with Omicron era variants benefitting the least (Figure 3D).

When interpreting these data, it is important to consider the temporal aspect of virus entry. Incoming PV’s encounter the exogenous, cell surface expressed TMPRSS2 first, and meet the endosomal cathepsins later, following endocytosis. For H655Y variants to show reduced sensitivity to TMPRSS2 overexpression, they must be somewhat resistant to proteolytic processing and activation at the cell surface. However, we have already established that this is unlikely to occur through changes in spike stability (Figure 2).

### Acquired interactions between H655Y and T696 do not dictate protease switching

We reasoned that novel intramolecular interactions, mediated by H655Y, may explain protease preference. First, we asked how prevalent Y655 is throughout the sarbecoviruses, allowing us to search for patterns in molecular interactions and structure. Drawing on sarbecovirus sequences from GenBank, we generated a maximum likelihood phylogenetic tree containing 278 spike sequences (Figure 4A). This revealed a cluster of spikes from bat-coronaviruses containing an equivalent H655Y, and some sporadic acquisitions of H655Y in SARS-CoV-1 sequences. The overall picture would suggest that H655Y is not a common occurrence, however the presence of a distinct lineage of H655Y bat coronaviruses, which lack a S1/S2 FCS, suggests that H655Y may not solely be a specific adaptation to furin cleavage as observed with SARS-CoV-2.

**Figure 4:**
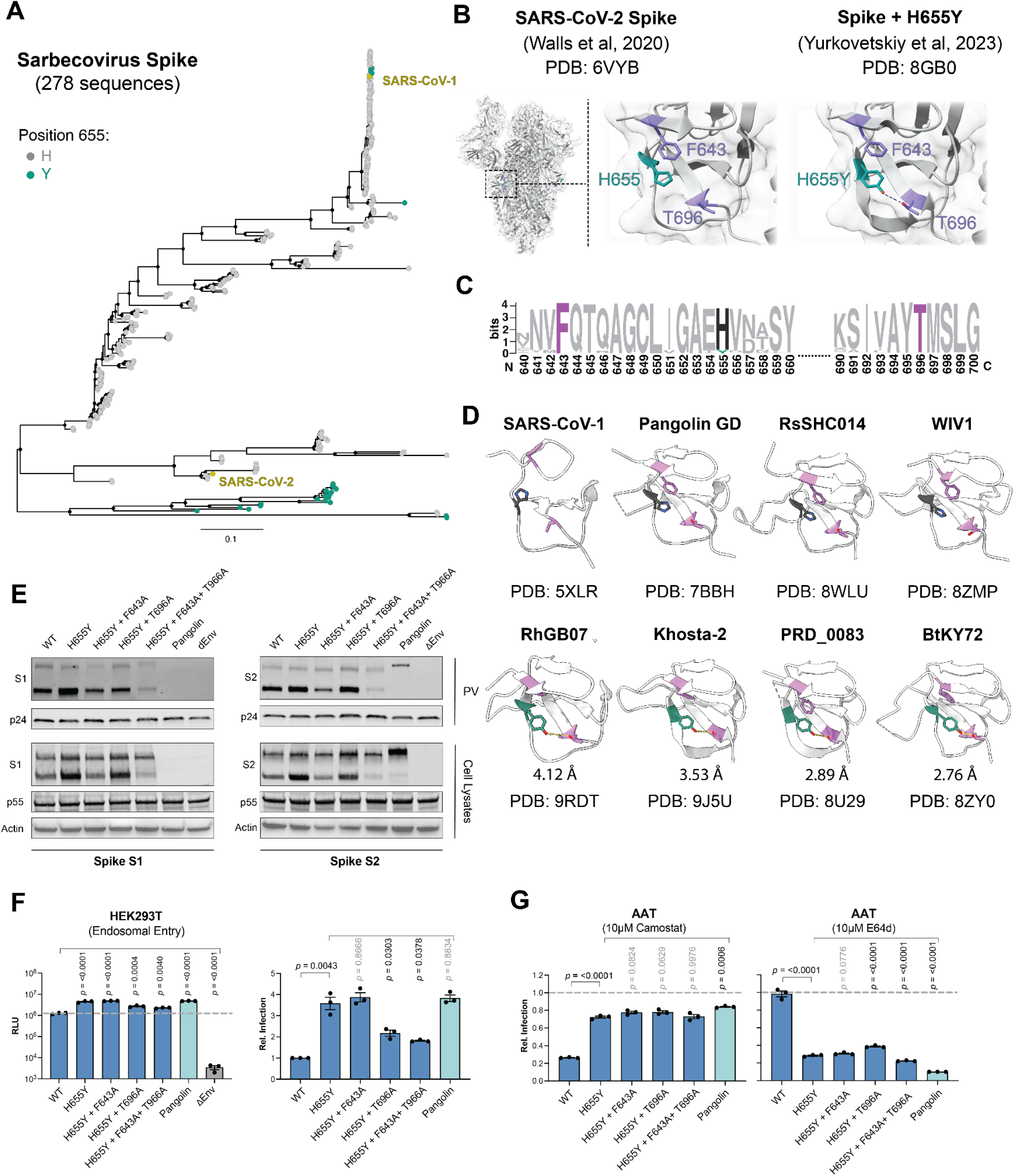
Evolution of H655Y in sarbecoviruses. **A.** Phylogenetic tree of 278 sarbecovirus spike sequences. Grey dots denote SARS-CoV-2 spike H655 equivalent residues, whereas green dots represent Y655. **B.** Zoomed-in structures of WT and WT+H655Y Spikes showing H655Y interactions with positions 643 and 696. **C.** Conservation plot of Spike residues from SARS-CoV-2 equivalent residue positions 640-700. **D.** PDB structures of sarbecovirus spikes that have either H655 or Y655 residues. Y655 to T696 distances were measured in 655Y containing viruses. **E.** Western blot panels of SARS-CoV-2 spike mutants. Lysates and PVs were immunoblotted against S1 and S2 (spike), p55/p24 (gag-pol control) and actin. **F.** Infectivity of mutant PVs in TMPRSS2-deficient HEK293T cells. dEnv is bald PV control. **G.** Sensitivity panel of mutant PVs in AATs in the presence of 10μM camostat or 10μM E64d. The dotted line represents relative viral infection with DMSO control. For **F** and **G,** statistical significance was determined by one-way ANOVA test was used against the mean of WT. *P* values ≤ 0.05 are considered significant. A minimum of n=3 was used for each comparison.

Yurkovetskiy et al. [43] resolved the structure of SARS-CoV-2 H655Y spike and showed an, acquired, polar interactions between H665Y, in S1, and T696, in S2 (Figure 4B). This interaction occurs in addition to pi-stacking (non-covalent interactions of the aromatic rings) within residue 655 and F643. Sarbecovirus spike alignment shows that F643 and T696 are highly conserved (Figure 4C). Comparison of structures of bat coronaviruses that contain either histidine or tyrosine at position 655 revealed a difference in local architecture driven by the formation of a polar interaction between Y655 and T696, in agreement with the structure from Yurkovetskiy et al (Figure 4D). We hypothesised that this interaction in SARS-CoV-2 variants containing H655Y may contribute to protease preference. To test this, we mutated either F643 (which participates in pi-stacking with H655Y), T696A (which mediates the novel polar interaction) or both in a H655Y background, expecting to revert protease preference to TMPRSS2.

First we assessed spike expression, particle incorporation and integrity by western blotting. T696A showed modest effects, with similar S1 and S2 levels in the PV fraction and cell lysate portion (Figure 4E). In contrast, F643A reduces both S1 and S2 levels in the cell lysate and PV fraction, suggesting that F643 is important for spike stability. The double mutant, F643A + T696A, had a compounding effect, leading to a pronounced decrease in S1 and S2, again suggesting a destablised spike protein.

To evaluate protease preference, we first used HEK293T cells, which are only permissive to endosomal entry (Figure 4F). We observed little correlation between spike incorporation in the PV and infection of HEK293T cells. For example, H655Y + F643A + T696A exhibited robust infection (∼two-fold greater than WT) despite reduced levels of spike. As expected, H655Y exhibited enhanced entry compared to WT. F643A did not affect cell entry compared to H655Y, whereas both T696A and the double mutant exhibited reduced infectivity, albeit whilst retaining a two-fold increase over WT spike. Taken alone, this result may suggest a change in protease preference, however, we could not yet distinguish generally reduced T696A spike activity from a change in protease preference. Therefore, we infected AAT cells with each PV, using camostat and E64d to differentiate entry (Figure 4G). This revealed there was no phenotypic switch upon mutation of either F643 or T696. This demonstrates that our hypothesis was incorrect and that molecular interactions between H655Y and these conserved residues do not contribute to protease preference.

### Mutations in NTD and RBD of BA.2.86 can override the H655Y phenotype

In our and others recent work [42,46], and across multiple experiments in this work (Figure 1B, 2B & G), the Omicron variant BA.2.86 consistently exhibits a more pre-Omicron-like phenotype; it is able to infect lung cells and is more sensitive to camostat inhibition, indicating a regained ability to use TMPRSS2. BA.2.86 possesses >30 mutations relative to its nearest ancestor, BA2, but retains H655Y. Given that, in all other genetic backgrounds, H655Y is a dominant mutation that controls activating protease usage (Fig. 1-3), we reasoned that BA.2.86 must possess overriding mutations that suppress the H655Y phenotype. To explore this, we performed further domain swaps, where we delineate both the NTD and RBD into two further splits, NTD1,2 and RBD1,2 (Figure 5A, 5B). We placed these ‘subdomain’ swaps of BA.2.86 in a WT and WT+H655Y background, and the reciprocal WT subdomains into a BA.2.86 backbone. Using AAT cells in the presence of protease inhibitors, we found that swapping NTD1 and RBD1 into the H655Y background renders spike camostat sensitive and E64d resistant, consistent with a reversion in protease preference (Figure 5C, 2D). This correlates with the importance of BA.2.86 mutations S50L (found in NTD 1) and K356T (found in RBD 1) for efficient infection of lung cells [46], and suggests that these, and potentially other mutations in the NTD and RBD, are able to nullify the otherwise dominant effects of H655Y.

**Figure 5:**
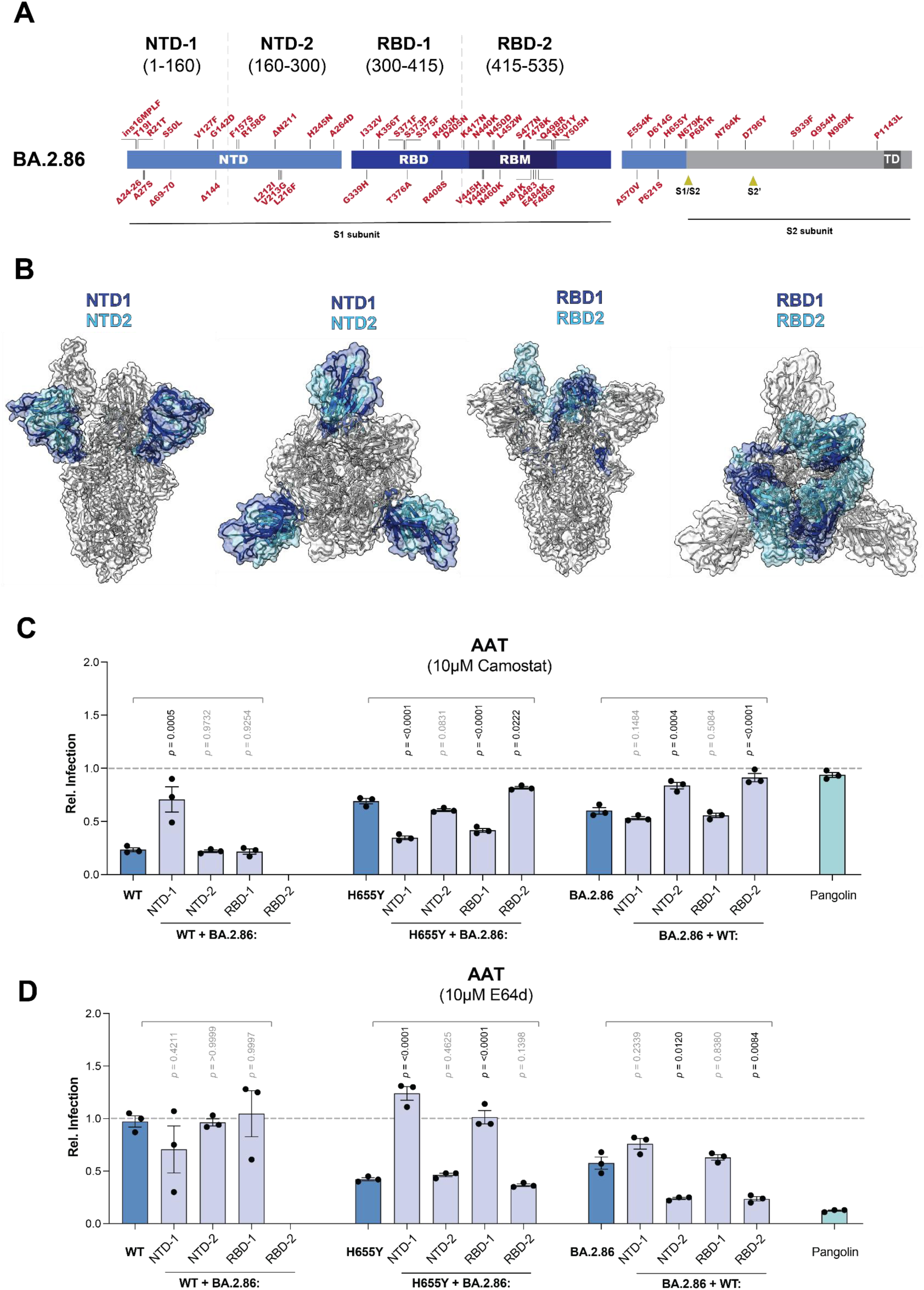
Subdomain swapping in BA.2.86. Ribbon diagram depicting the mutational profile of BA.2.86. Boundaries of each subdomain swap are denoted by a grey line. **B.** Structural depiction of subdomain swaps using WT spike structure. **C, D.** Sensitivity panel of subdomain swapped PVs in AATs in the presence of 10μM camostat or 10μM E64d. The dotted line represents relative viral infection with DMSO control. WT + BA.2.86 RBD2 swaps did not rescue. **D.** For **C** and **D,** statistical significance was determined by one-way ANOVA test was used against the mean of WT. *P* values ≤ 0.05 are considered significant. A minimum of n=3 was used for each comparison.

### H655Y is not a determinant of upper airway cell tropism

Thus far, we have demonstrated that H655Y is the genetic determinant of activating protease preference and, therefore, explains a defining *in vitro* phenotype of Omicron lineage viruses: endosomal entry. Next, we asked whether this was directly linked to another hallmark of Omicron viruses: enhanced infection of upper-respiratory cells. Using an authentic virus setting, we have recently demonstrated that Omicron defining mutations in the RBD influence cellular tropism [42], and we hypothesised that H655Y would be a necessary genetic background for this phenotype. We tested this with authentic virus in various cell systems, using reverse genetics to toggle the residue 655 genotype of WT and BA.1 viruses, and infected AAT cells, Calu3 cells and human nasal epithelial cells (hNEC).

We initially tested whether H655Y and its reversion in an authentic virus background is consistent with the phenotypes observed in PV. Our AAT infection assay in the presence of drug inhibitors demonstrated that H655Y unlocked cathepsin usage in WT virus, therefore recapitulating our PV infection assays (Figure 6A). However, BA.1 + Y655H only moderately re-engaged with TMPRSS2-mediated entry. It was somewhat sensitised to camostat and resistant to E64d. This effect was not as dramatic as observed in PV, but is consistent with our previous work in which BA.1 authentic virus exhibits a more subtle entry phenotype compared to contemporary Omicron viruses such as BA.2 [18,42].

**Figure 6:**
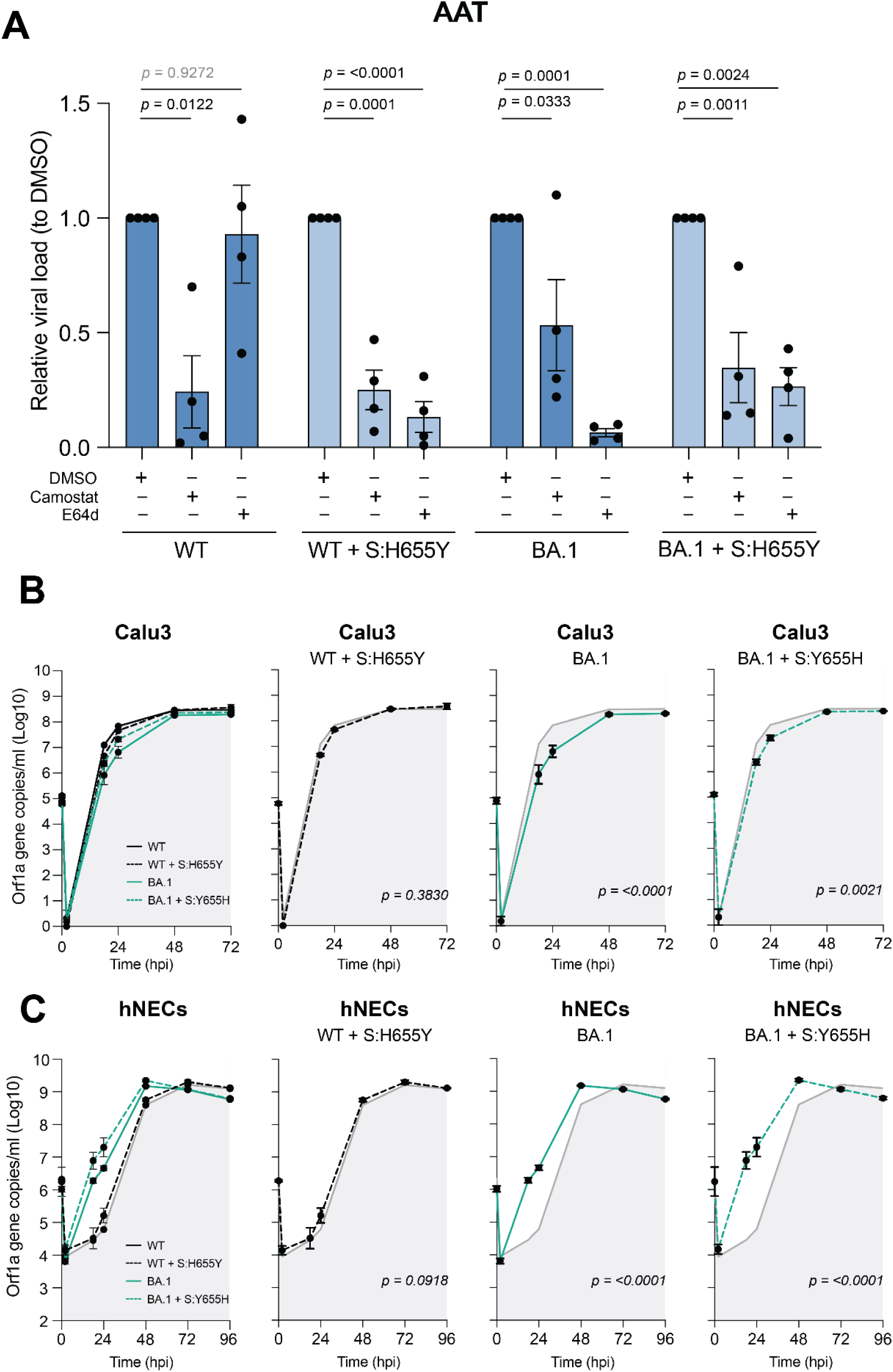
Authentic virus upper airway infections. Sensitivity panel of authentic virus mutants in AAT cells in the presence of 10μM camostat or 10μM E64d. **B.** Replication kinetics of authentic viruses in Calu3 cells quantified by RT-qPCR at different timepoints post infection (hpi) as indicated. SARS-CoV-1 B.1 was used as a reference virus in each experiment (as denoted by grey area). Note that values at time 0 hpi reflect the tirtres of input virus, whilst those at 1 hpi are titres of residual virus in the supernatant after an 1 hour post infection and one wash of the monolayer. **C.** Replication kinetics of authentic viruses in primary nasal epithelium cells quantified by RT-qPCR at different timepoints post infection (hpi) as indicated. SARS-CoV-2 B.1 was used as a reference virus in each experiment (as denoted by the grey area). For **A, B** and **C,** statistical significance was determined by one-way ANOVA test were used to determine significance. *P* values ≤ 0.05 are considered significant.

Calu3 cells are lung-derived and express high levels of TMPRSS2 and, therefore, act as a surrogate for lower-airway cells. Whilst we observed a decrease in infection and replication with BA.1 (Figure 6B), confirming its preference for upper airway cells [42], introduction of H655Y in WT did not affect viral replication in Calu3 cells. Y655H did somewhat rescue BA.1 replication at the 24 hrs timepoint suggesting that this mutation makes a modest contribution to impaired lower-airway tropism in an Omicron genetic background.

The critical experiment however was to assess the importance of H655Y for enhanced replication in hNAEC, a key characteristic of Omicron lineage viruses. Consistent with our and others previous work, we observed enhanced replication of BA.1 in hNEC, compared to WT (Figure 6C). Notably, WT+H655Y or BA.1+Y655H did not impact on viral replication in hNEC, demonstrating that H655Y is neither necessary nor sufficient for upper airway tropism. This somewhat surprising result suggests that two of the defining characteristics of Omicron, a switch in activating protease and a preference for upper-airway replication are distinct phenotypes that are separable through a single point mutation at spike residue 655.

## Discussion

SARS-CoV-2’s emergence with an FCS marked an evolutionary departure from other sarbecoviruses, enabling spike pre-processing for rapid cell surface fusion. The emergence of the Omicron lineage heralded two pronounced phenotypic changes in SARS-CoV-2: an apparent switch in preference for spike-activating protease, and a change in tissue tropism preference towards the upper respiratory tract. The extent to which these features were mechanistically linked was unclear. Whilst initial hypotheses focused on impaired FCS processing [17], our data and other studies have confirmed efficient cleavage of spike on the surface of particles, distinct from cell lysates, indicating that mutations beyond the FCS drive this switch. Therefore, in this study, we aimed to determine the genetic and molecular determinants of protease preference and to understand how this relates to tissue tropism.

We, amongst others, confirmed that H655Y drives the switch from TMPRSS2-mediated cell surface entry to cathepsin-dependent endosomal entry [19,20,22]. This effect is two-fold, whereby H655Y both rescues endosomal entry but also reduces spike sensitivity to TMPRSS2. A leading hypothesis proposed that H655Y increases spike stability [19,43], thereby limiting cell surface entry by reducing S1 shedding: a pre-requisite event for membrane fusion following binding to ACE2 [47–49]. This mechanism and hypothesis was consistent with our observations and seemed plausible given that our CMK-mediated prevention of furin processing effectively reverted all the variants to endosomal entry. Previous work on SARS-CoV-1 also demonstrated that exogenous processing of S1/S2, which permits early shedding of S1, promotes cell surface entry [50]. However, western blots with our N-terminal HA-tagged spike variants western blots dispute this hypothesis on several grounds. Whilst S1 levels were subtly higher in WT+H655Y, variants D614G and Alpha showed the highest S1 incorporation despite lacking H655Y, yet readily utilised cell surface entry. Conversely, many of the Omicron variants showed reduced retention of S1, yet enter endosomally. Furthermore, F643 and T6969 residues, which were proposed to strengthen S1/S2 interactions via H655Y [43], showed variable S1 levels by western blot and all retained the same endosomal entry phenotype, despite their substitution to alanines. Taken together our findings indicate no correlation between S1 retention and protease preference, suggesting that S1 shedding is not the primary mechanism underlying the H655Y phenotype.

Nonetheless, very recent observations allow us to consider other mechanistic explanations. Structural analysis indicates that H655 is solvent exposed, a property lost upon mutation to Y655 [43]. Given H655Y’s proximity to the neighbouring monomer’s S2’ cleavage site, a target for TMPRSS2 processing at the cell surface, we speculate that H655Y directly modulates S2’ cleavage efficiency. This interpretation aligns with another report [51] that observed a delay in entry when H655Y is present, attributed to stabilisation of an intermediate state of spike, thus altering the membrane fusion kinetics. We find this model interesting, as a subsequent structural study suggests that TMPRSS2 cleavage of S2’ can only occur on ACE2-bound spikes that have adopted an intermediate conformation between pre- and post-fusion, subsequently leading to S1 shedding and membrane fusion [52]. Furthermore, the same study reports that the S2’ recognition sequence is buried when spike is in a pre-fusion ACE2-bound state, only to be exposed in the intermediate state. Building upon Yurkovetsky et al [43], Qing et al [51] and McCallum et al [52], we proposed that H655Y may modulate the kinetics of conformational transitsions such that early activation by TMPRSS2 is disfavoured, allowing particles to undergo endocytosis and complete activation in the endosome. Ultimately, confirmation of this, and a definitive mechanistic explanation, would require determination of co-structures of TMPRSS2 with spike, and an understanding of how H655Y directly alters the availability of S2’ during entry.

Despite the incomplete mechanistic understanding of the H655Y phenotype, we reasoned that this mutation would also be necessary for the switch in tissue tropism. Our assays using primary upper airway respiratory cells have demonstrated conclusively that neither H655Y nor its reversion in a BA.1 background were insufficient for reverting tropism, thus delineating protease preference and tissue tropism. In our previous work [42], we identified a cluster of Omicron RBD mutations, D339, F371, P373 and F375, as strong drivers of tropism switch from lower lung to upper airway, though substitution of those residues in a WT (B.1) background were insufficient to recapitulate infection in hNECs. Therefore, the precise mechanism is still unknown. Several explanations have been proposed, such as acquisition or enhanced interactions with alternate matrix metalloproteinases [29] (MMPs), disintegrin and metalloproteinases [30] (ADAMs) family of proteases, heparan sulfate [53] and sialic acids [31]. In any case, we can state that tropism switch is complex and not attributable to a singular mutation acting as an on/off switch for protease usage, and appear to work in context. Further work is required to understand what other factors are involved, and how mutations in the RBD determine tropism.

In summary, we show that, in most genetic backgrounds, H655Y is a dominant determinant of protease preference, irrespective of pre-processing status at the FCS. Moreover, H655Y simultaneously rescues endosomal entry, via cathepsins, and reduces sensitivity to TMPRSS2-mediated activation. Whilst a complete mechanistic understanding has not been forthcoming, recent observations, by others, suggest that changes to the dynamics and choreography of the spike fusion mechanism may underpin the H655Y phenotype. Critically, we also demonstrate that protease usage and enhanced nasal epithelial cell infection represent distinct and separable phenotypes. This discovery will be valuable in distentangling the complex determinants of SARS-CoV-2 tissue tropism and, by extension, the drivers of virus transmission and epidemiology.

## Supporting information

Supplementary Western Blot Files

Supplementary Raw Datasets

## Acknowledgements

This work was funded by UK Medical Research Council (MRC) support to the MRC-University of Glasgow Centre for Virus Research (CVR Structure-to-Function of Virions Programme: MC_UU_00034/1 and CVR Preparedness Platform: MC_UU_00034/6, to J.G., M.P., A.H.P., B.J.W.). We received additional support from the MRC to the G2P2 consortium (MR/Y004205; to M.P. and A.H.P.), and the Wellcome Trust to the G2P-Global consortium (226141/Z/22/Z; to M.P.). The funders had no role in study design, data collection and analysis, decision to publish or preparation of the manuscript.

