## Supplementary Western Blot Files for "SARS-CoV-2 Spike H655Y drives protease preference but does not dictate cellular tropism"

### Raw blot images used for Figure 2C.

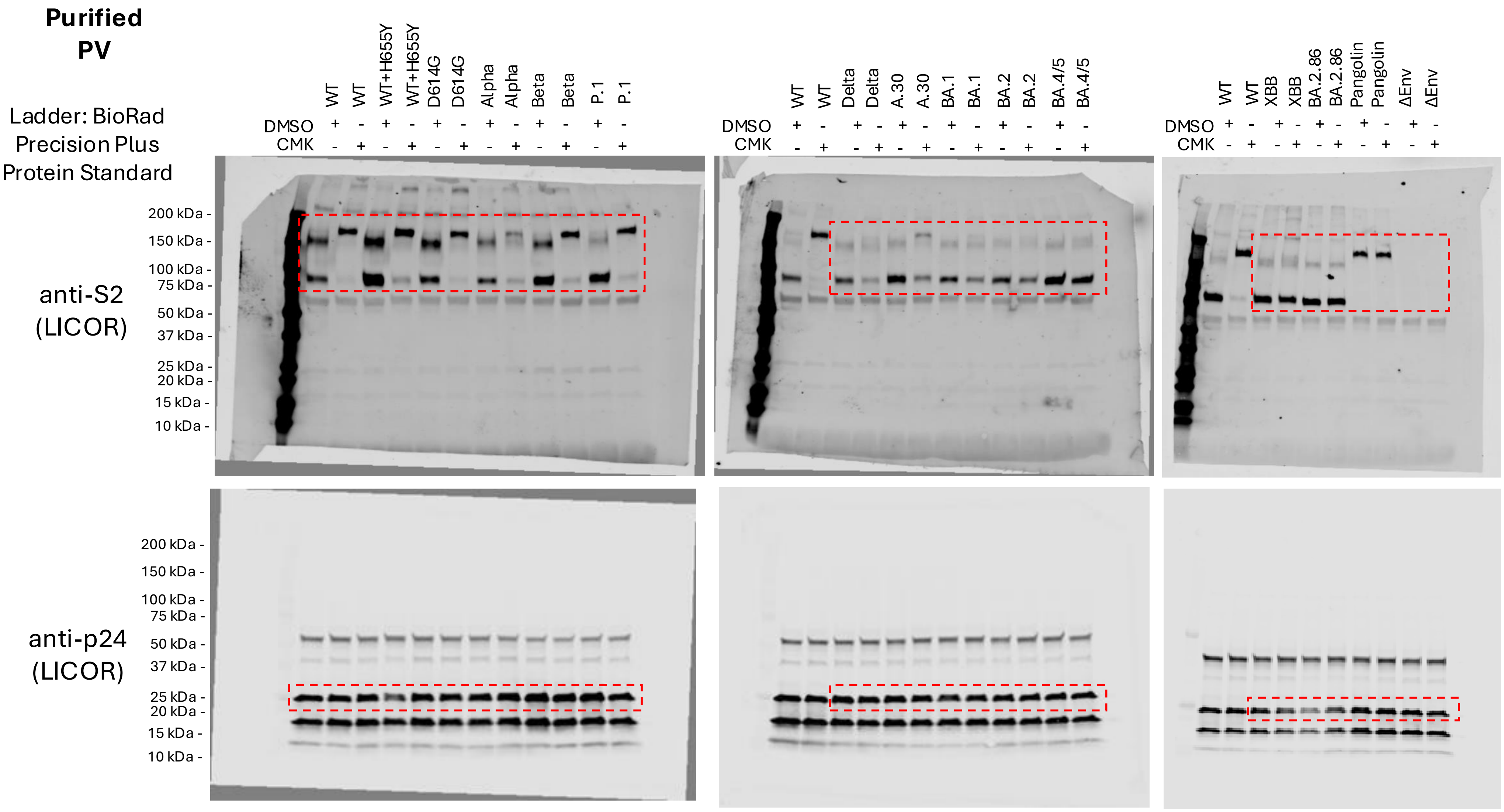

### Raw blot images used for Figure 2C.

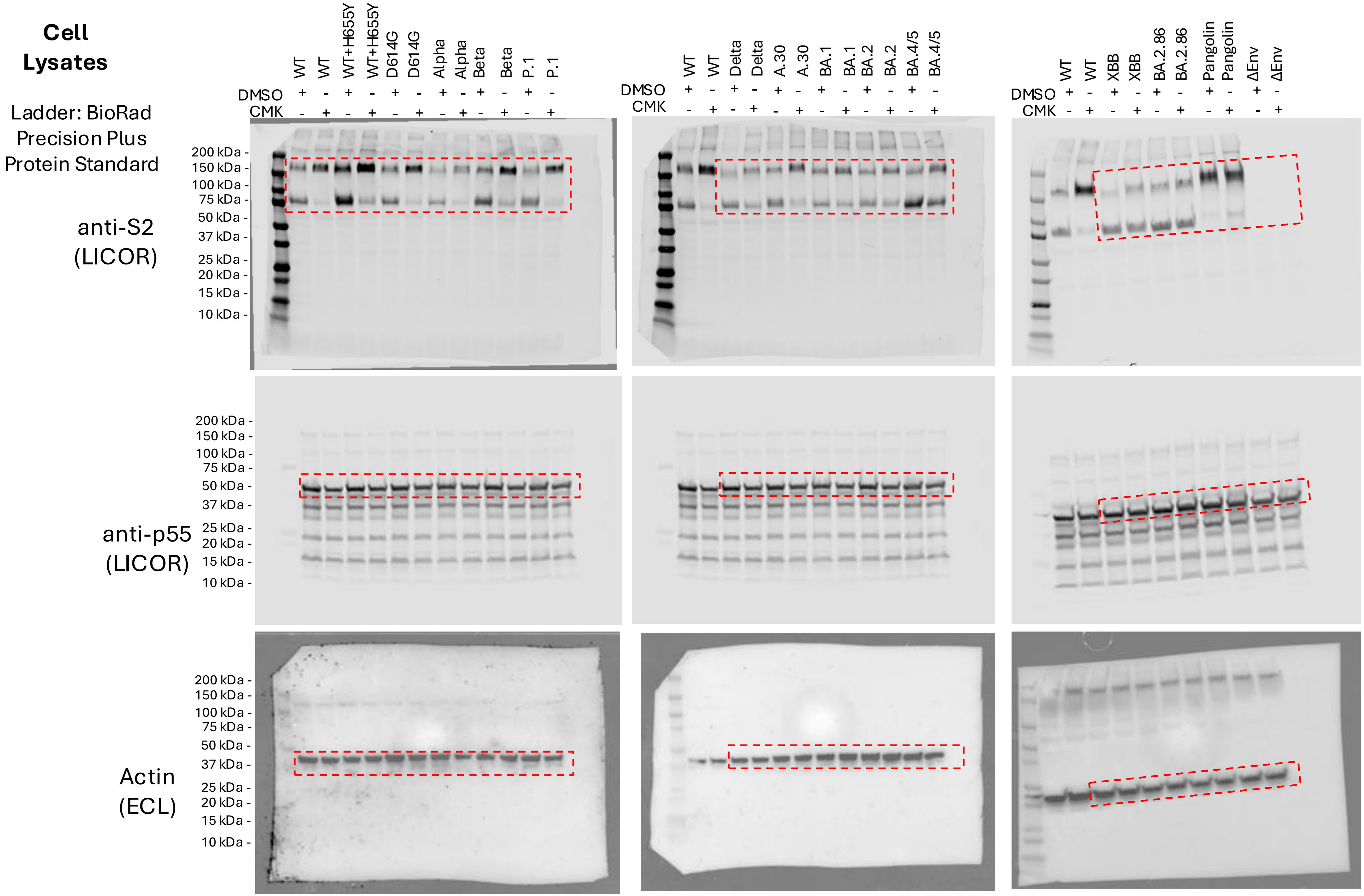

### Raw blot images used for Figure 2H

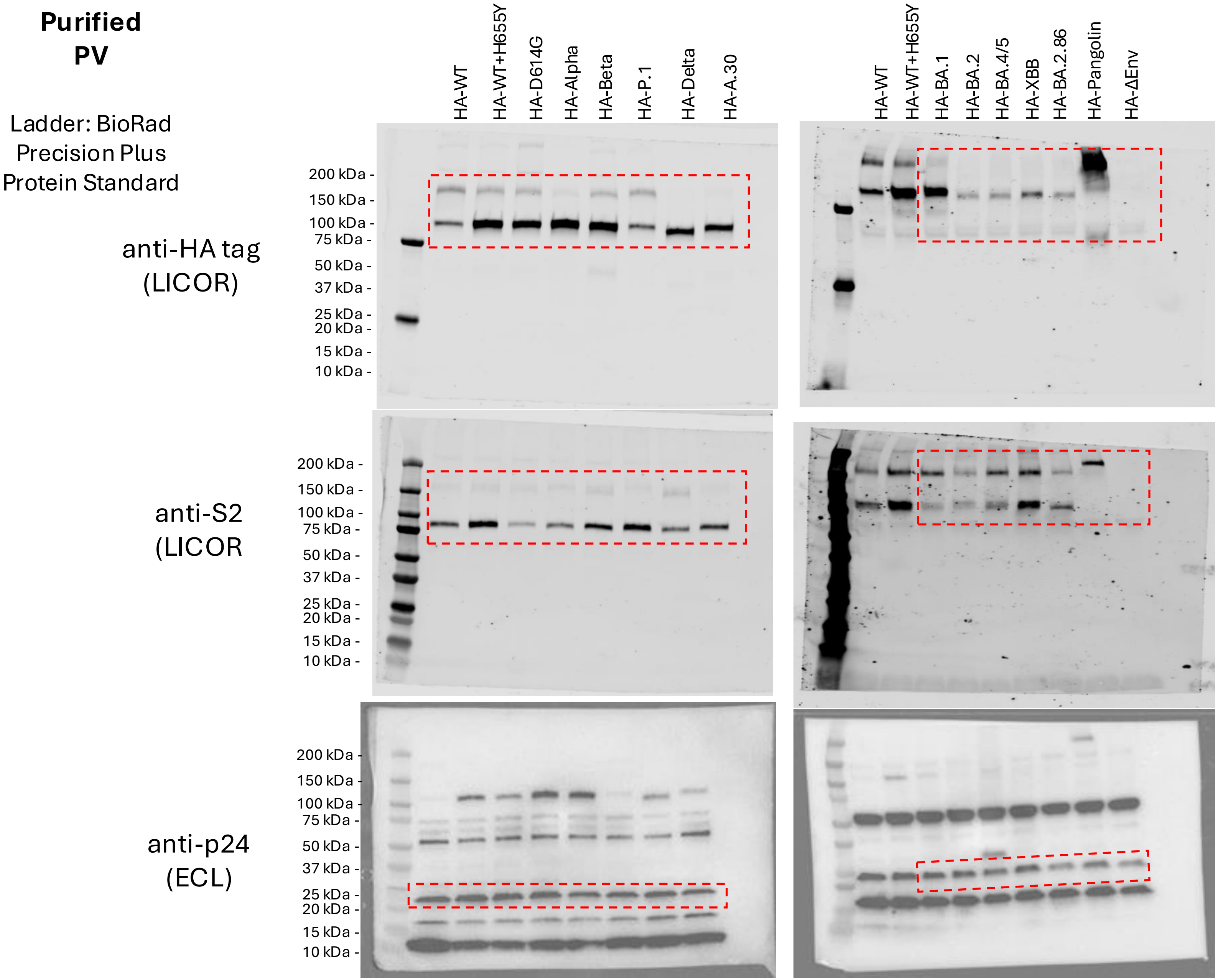

### Raw blot images used for Figure 2H

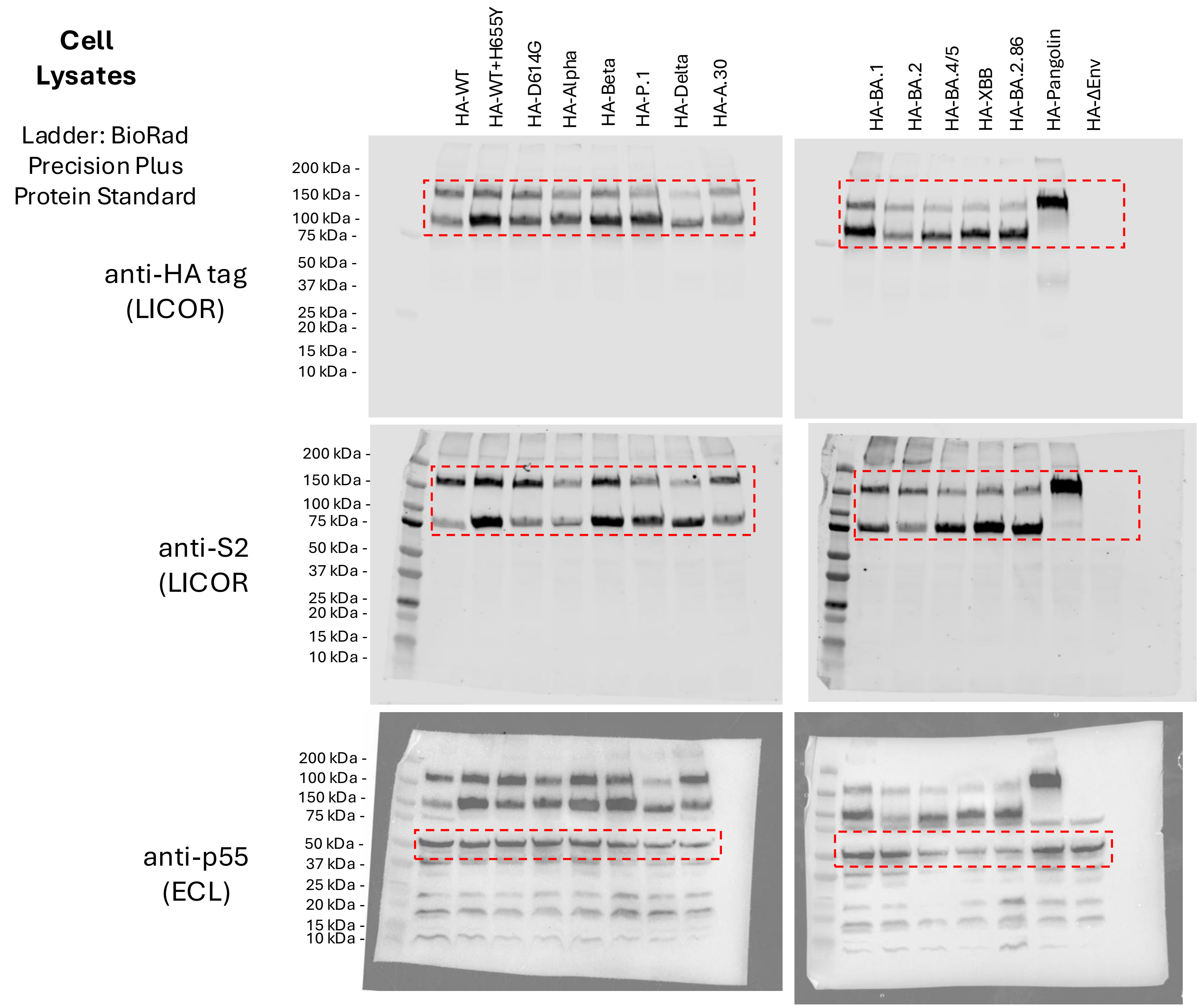

### Raw blot images used for Figure 4E

**Purified PV**

Ladder: BioRad Precision Plus Protein Standard

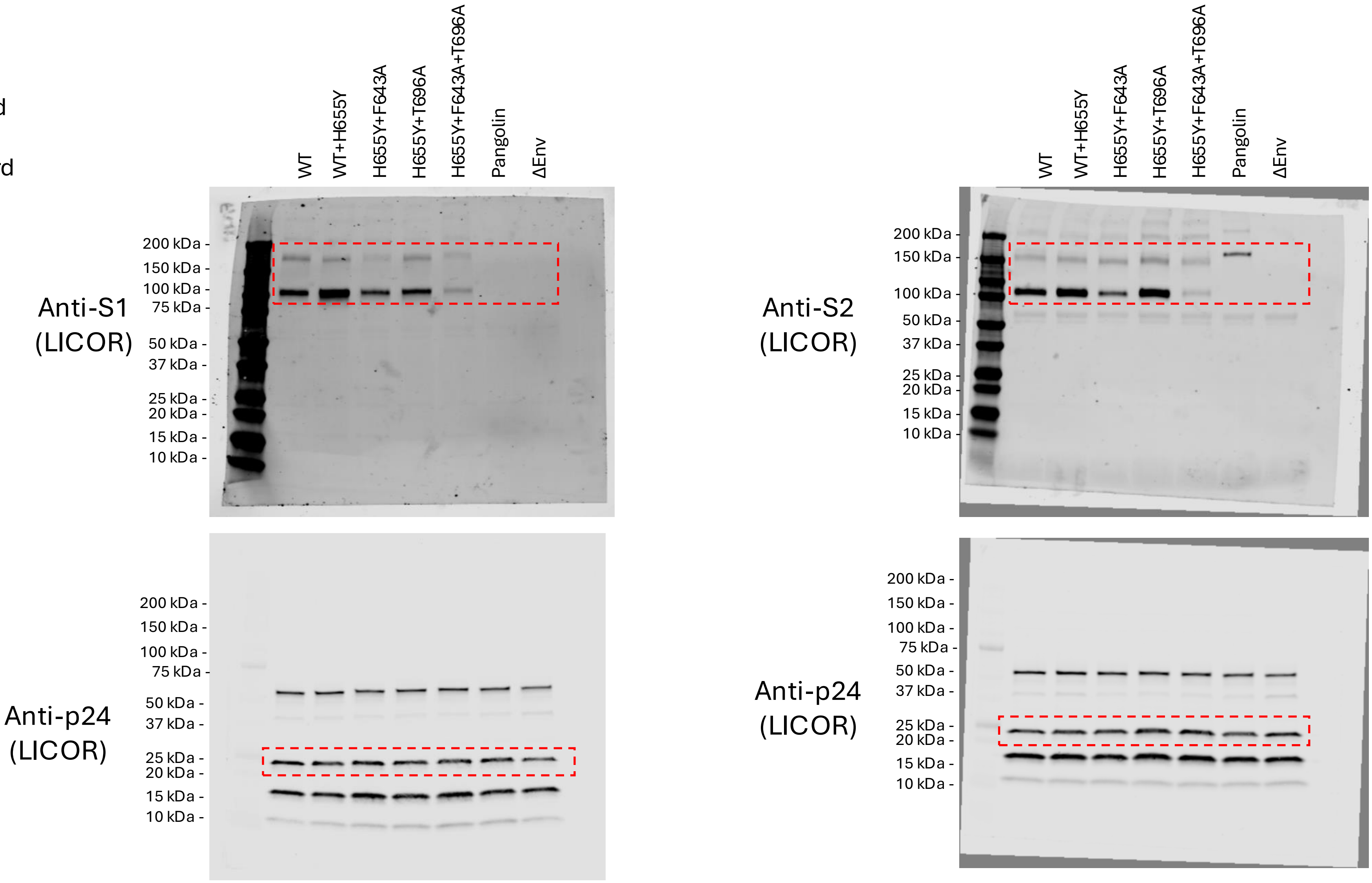

### Raw blot images used for Figure 4E

**Cell Lysates**

Ladder: BioRad Precision Plus Protein Standard

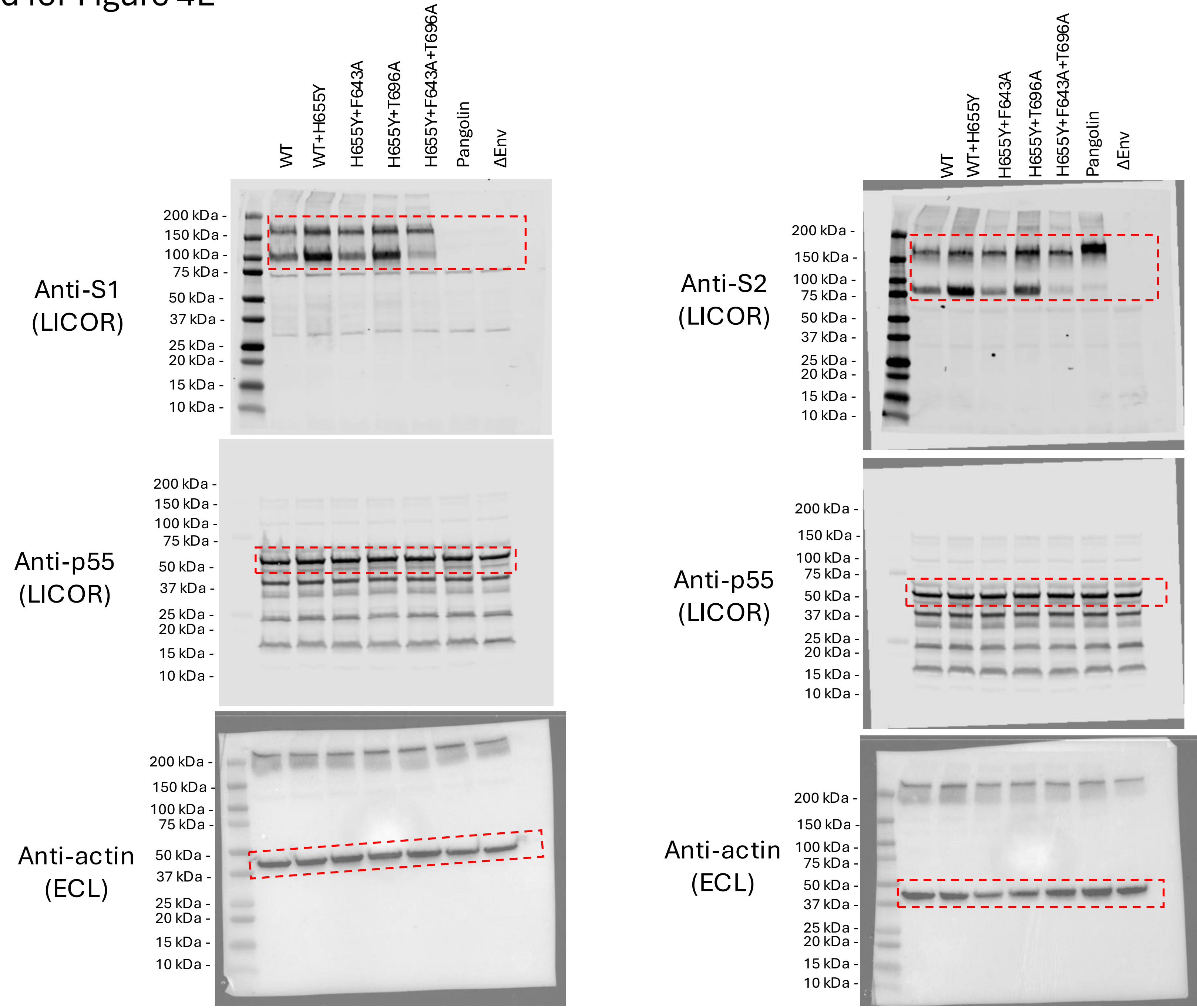
